# Integrated single-cell sequencing analysis reveals peripheral immune landscape across the human lifespan

**DOI:** 10.1101/2022.07.11.498621

**Authors:** Renyang Tong, Ronghong Li, Yichao Zhao, Lingjie Luo, Jianmei Zhong, Zheng Li, Lin Wei, Yifan Chen, Jianfeng Shi, Yu Gao, Mingze Sun, Yuyan Lyu, Ancai Yuan, Lingyue Sun, Yuping Guo, Honglin Wang, Haoyan Chen, Lei Chen, Bin Li, Wei Lin, Feng Wang, Li Wang, Jun Pu

## Abstract

Systematic understanding of immune dynamics across the entire human lifespan at single-cell resolution is currently lacking. Here, we performed single-cell RNA sequencing (scRNA-seq) and single-cell T cell receptor (TCR)/B cell receptor (BCR) sequencing (scTCR/BCR-seq) on over 380,000 peripheral blood mononuclear cells collected from 45 healthy participants aged 0 to over 90 years. We revealed that the functions of T cell subsets were most susceptible to senescence among all PBMCs, featured by increased NF-κB signaling and IFN-γ responses pathways, while reduced telomere maintenance and energy metabolism. By combined analysis of scRNA-seq and scTCR-seq, we revealed that 1) Naïve CD4^+^ T and naïve CD8^+^ T cells displayed different aging patterns in both transcriptomes and immune repertoires; and 2) CD8^+^ MAIT cells showed a higher cell abundance and clonal diversity in adolescents. In summary, our work provided valuable insights and rich resources for understanding the development and aging of the human immune system across the lifespan.

## INTRODUCTION

Development and aging run through the human life with gradual maturation and inevitable degeneration of the immune system (Mittelbrunn and Kroemer, 2021; Vijg and Calder, 2004). As a key defense system, immune system protects the host against various exogenous invasions and endogenous challenges (e.g., viruses, bacteria, parasites, and cancer cells) (López-Otín and Kroemer, 2021). Human beings tend to be less vulnerable to foreign pathogens as the immune system gradually matures after birth (Simon et al., 2015). By contrast, immunosenescence leads to a higher incidence of infection, autoimmune diseases and cancer with aging (Montecino-Rodriguez et al., 2013; Nikolich-Zugich, 2018). Moreover, aged immune cells drive aging of solid organs, such as the heart and liver, which is a probable cause of systemic senescence (Yousefzadeh et al., 2021). Yet, little is known about when and how the immune system changes across the entire lifespan. Therefore, there is an urgent need to systematically understand the physiology/pathology of immune development and immune senescence across the human lifespan.

Human peripheral blood mononuclear cells (PBMCs) are a diverse mixture of highly specialized immune cells that are critical components of the immune system. Prior omics studies of human PBMCs have revealed their involvement in immune pathologies [e.g., immunosenescence (Alpert et al., 2019; Mogilenko et al., 2021; Zhang et al., 2019), autoimmune disorders (Nehar-Belaid et al., 2020) and infectious diseases (Ren et al., 2021; Zhu et al., 2020)] as well as in several diseases of solid organs, such as cardio-cerebrovascular diseases (Mazzola et al., 2008; Moore et al., 2005) and acute-on-chronic liver failure (Li et al., 2022). Some significant age-related changes in PBMCs composition and function from healthy donors have been demonstrated using flow cytometry or bulk sequencing. For example, flow cytometry analysis demonstrated that numbers of CD19^+^ B cells and naive T cells decreased with age, whereas antigen-experienced memory T cell numbers increased with concomitant loss of co-stimulation factors CD27 and CD28 (Fagnoni et al., 2000; Qin et al., 2016). Moreover, bulk RNA sequencing identified a large group of age-associated genes in PBMCs and whole blood, which was used to construct prediction models for “transcriptomic age” (Peters et al., 2015). These findings provided valuable insights into the mechanism of immune aging, whereas these assays based on canonical cell markers and classification criteria might be insufficient to reflect the sophistication and heterogeneity of peripheral immune cells.

Single-cell RNA sequencing (scRNA-seq) and single-cell T cell receptor (TCR)/B cell receptor (BCR) sequencing (scTCR/BCR-seq) provide an opportunity to comprehensively identify previously undefined cell types at single-cell resolution (Ahmed et al., 2019; See et al., 2017; Villani et al., 2017). It can provide novel details into both development- and aging-related processes by deconvoluting molecular features within specific cell types (Han et al., 2020; Ma et al., 2020). Several scRNA-seq studies on peripheral immune cells conducted at specific ages have yielded valuable findings at single-cell resolution: 1) compared with adult PBMCs from public database, scRNA-seq of neonate umbilical cord blood (UCB) identified an enrichment of a previously unrecognized cytotoxic innate immune cell type, namely GZMB^+^ natural killer T (NKT) cells, although the specific functions of this UCB-specific immune cells remain unclear (Zhao et al., 2019); 2) compared with young controls (25 to 29 years old), clonal GZMK^+^ CD8^+^ T cells were identified as a hallmark of inflammaging in order individuals (62 to 70 years old) (Mogilenko *et al*., 2021); and 3) compared with senior controls (aged 50 to 89 years old), scRNA-seq data revealed the expansion of cytotoxic CD4^+^ T cells as a key feature of healthy immunosenescence in very elderly individuals (i.e., supercentenarian reaching 110 years of age) (Hashimoto et al., 2019). These scRNA-seq studies comparing between two specific age groups provided valuable insights into age-related changes in peripheral immune cells at single-cell resolution. However, a systematic understanding of peripheral immune cell dynamics across the entire human lifespan at single-cell resolution is currently lacking.

Here, by combined analysis of scRNA-seq and scTCR/BCR-seq, we systematically investigated dynamic changes in the cell-type composition, transcription profiles, and immune repertoires of human peripheral immune cells across the entire lifespan, using PBMC samples from 45 healthy participants aged 0 (neonates) to ≥ 90 y (nonagenarians). Our work provided a rich resource for understanding the development and aging of human peripheral immune cells. We compiled these analyses into a web portal (https://pu-lab.sjtu.edu.cn/shiny) for exploring the data easily.

## RESULTS

### Overview of single-cell transcriptomes of peripheral immune cells across the entire lifespan

To characterize the dynamics of peripheral immune cell composition and function throughout the human lifespan, 45 healthy participants aged 0 (neonates) to ≥ 90 years (nonagenarians) were recruited, which spanned ten age groups including five neonates (umbilical cord blood, UCB group), three infants aged 1 year (1 y group), five toddlers aged years (2 y group), six preadolescents aged 6 years (6 y group), four adolescents aged 12 years (12 y group), five adolescents aged 18 years (18 y group), four young adults aged 30 years (30 y group), five middle-aged adults aged 50 years (50 y group), five septuagenarians aged 70 years (70 y group), and four nonagenarians aged ≥ 90 years (90 y group). All participant details are shown in Table S1. We performed single-cell RNA, TCR and BCR sequencing simultaneously on PBMCs from each individual, investigated the dynamics of cell type composition and immune cell functions across the lifespan (Figure 1A).

**Figure 1.**
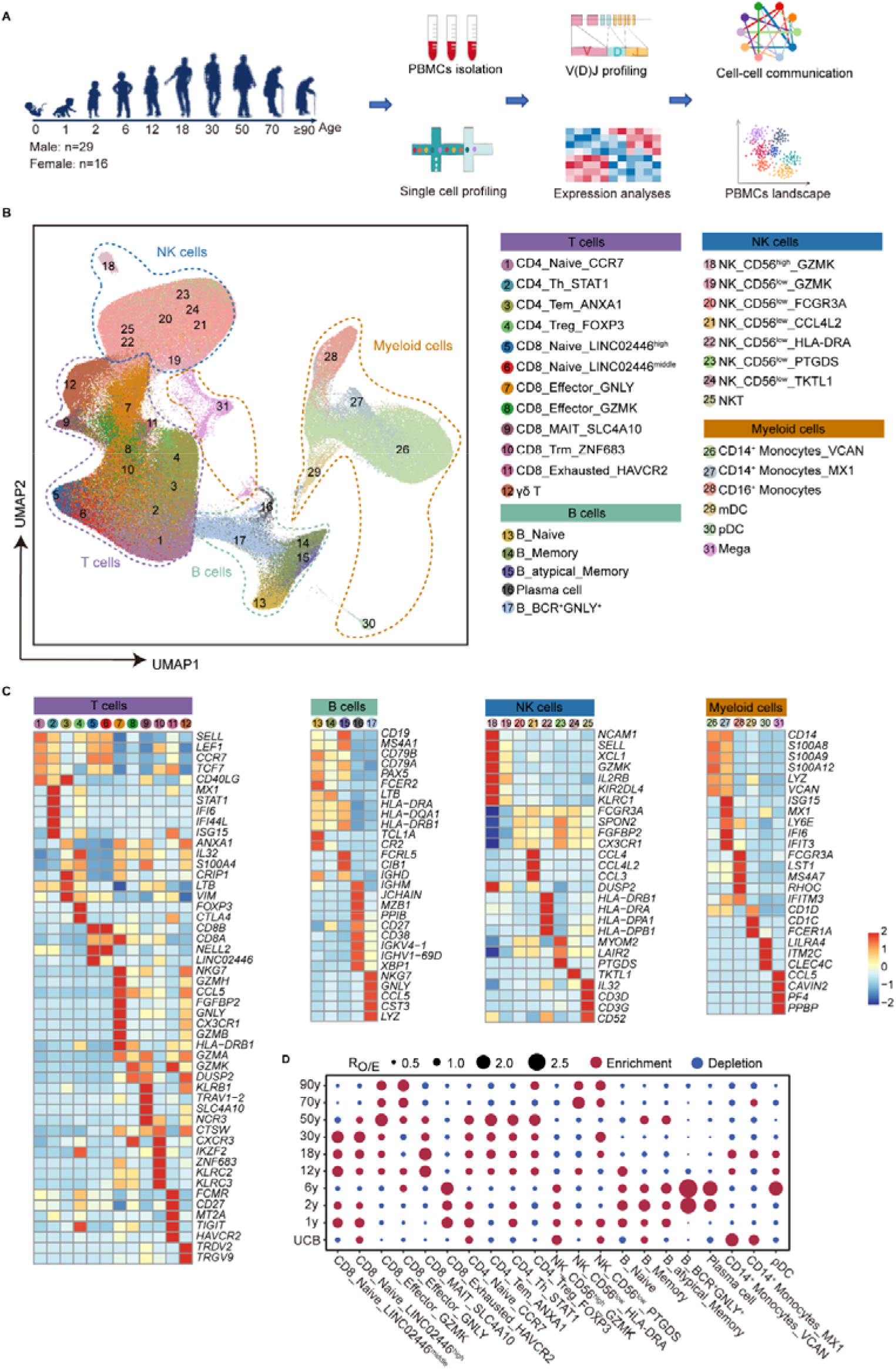
Overview of the single-cell transcriptome of peripheral immune cells across the entire lifespan. (A) Workflow of the overall experimental design of this study. (B) UMAP charts of more than 380,000 single immune cells from 45 human individuals. In the UMAP, 31 immune cell subsets are labelled with different colors. (C) Heatmap showing the expression of signature genes in each cell subset. Red, high expression; blue, low expression. (D) Age group preference of each cell subset measured by the ratio of observed to randomly expected cell numbers (R_O/E_). mDC, myeloid dendritic cells; pDC, plasmacytoid dendritic cells; Mega, megakaryocytes.

After filtering out low-quality droplets, more than 380,000 cells were obtained from all samples. The filtered data were integrated, dimension reduced and clustered in an unsupervised method. Based on the unsupervised clustering and expression of signature genes for each subset (Table S2), 31 distinct PBMC subsets were identified and visualized via uniform manifold approximation and projection (UMAP) (Figures 1B and S1A). Specifically, there were four CD4^+^ T cell subsets, including naïve CD4^+^ T cells (CD4_Naive_CCR7), effector memory CD4^+^ T cells (CD4_Tem_ANXA1), type 1 helper CD4^+^ T cells (CD4_Th_STAT1), and regulatory CD4^+^ T cells (CD4_Treg_FOXP3); seven CD8^+^ T cell subsets, including two types of naïve CD8^+^ T cells (CD8_Naive_LINC02446^middle^ and CD8_Naive_LINC02446^high^), two types of effector CD8^+^ T cells (CD8_Effector_GZMK and CD8_Effector_GNLY), tissue resident memory CD8^+^ T cells (CD8_Trm_ZNF683), mucosal-associated invariant CD8^+^ T cells (CD8_MAIT_SLC4A10) and exhausted CD8^+^ T cells (CD8_Exhausted_HAVCR2); one subset of γδ T cells; eight NK cell subsets, including NK cells with high expression of *NCAM1*(encoding CD56) (NK_CD56^high^_GZMK), and six NK cell subsets with low expression of *NCAM1* (NK_CD56^low^_GZMK, NK_CD56^low^_FCGR3A, NK_CD56^low^_CCL4L2, NK_CD56^low^_HLA-DRA, NK_CD56^low^_PTGDS and NK_CD56^low^_TKTL1), and NKT cells; five BCR-positive cell subsets, including naïve B cells (B_Naive), memory B cells (B_Memory), atypical memory B cells (B_atypical_Memory), plasma cells (Plasma cell), and a previous undefined, unique B cell subset (we named it as B_BCR^+^GNLY^+^, as detailed below); six myeloid cell subsets, including CD14^+^ classical monocytes (CD14^+^ Monocytes_VCAN), CD14^+^ monocytes with high expression of *MX1* (CD14^+^ Monocytes_MX1), CD16^+^ monocytes, myeloid dendritic cells (mDCs), plasmacytoid dendritic cells (pDCs), and megakaryocytes (Mega). The representative signature genes of all these cell subsets and their expression levels were shown in heatmaps and feature plots (Figures 1C and S1B).

Among BCR-positive cell subsets, we identified a previously unrecognized B cell type. We named this new-identified B cell type as B_BCR^+^GNLY^+^, since it highly expressed general genes encoding B cell markers (e.g., BCR and *JCHAIN*) and canonical cytotoxic genes such as *GNLY* and *NKG7*, but did not express several other canonical B cell marker genes (i.e., *MS4A1, CD19* and *CD79B*) (Figures 1C and S1B), indicating that these cells might represent a cytotoxic B cell type. Meanwhile, negative TCR, *TRAC* and *KLRF1* expression excluded the possibility of these cells being T and NK cells. Moreover, we excluded the potential technical artifact (i.e., multiplets) based on the following: 1) the number of detected genes in the B_BCR^+^GNLY^+^ cells were relatively lower than that in other cell subsets (Figure S1C); and 2) a stringent doublet filtration standard was performed to exclude potential doublets.

### Cell composition dynamics of peripheral immune cells across lifespan

To obtain a global view of the dynamics of PBMC composition throughout human life, the proportion of each of the 31 cell subsets in total PBMCs was calculated for each sample (Figures S2A). Among them, 21 cell subsets exhibited significant differences in their proportion in total PBMCs across different age groups (ANOVA, FDR < 0.05). To further illustrate the age preference of these cell subsets, the ratio of observed cell numbers to randomly expected ones (R_O/E)_) (Zhang et al., 2018) was calculated. All these 21 cell subsets exhibited differential age preference (Figure 1D). Among these subsets, the proportion of certain cell subsets was positively correlated with age, as exemplified by CD8_Effector_GZMK cells (Spearman correlation coefficient R = 0.7082, p = 5.26×10^−8^), CD4_Treg_FOXP3 cells (R = 0.5174, p = 0.0002) and NK_CD56^low^_HLA-DRA cells (R = 0.4882, p = 0.0006) (Figures 1D and S2B). By contrast, the proportion of certain cell subsets was negatively correlated with age, such as NK_CD56^high^_GZMK cells (R = -0.5936, p = 1.72×10^−5^) and B_Naive cells (R = -0.5579, p = 6.83×10^−5^) (Figures 1D and S2C). Several cell subsets exhibited an enrichment at certain age stages. For example, CD14^+^ Monocytes_VCAN cells were enriched in UCB group; B_BCR^+^GNLY^+^ and plasma cells were enriched in children (2 y and 6 y groups); and CD8_MAIT_SLC4A10 cells were enriched in adolescents (12 y and 18 y groups) (Figures 1D, and S2D). Overall, we detected differential correlations of distinct PBMC subsets with age at single-cell resolution.

### Global transcriptome dynamics of peripheral immune cells throughout lifespan

To systematically investigate the transcriptome dynamics of PBMC subsets across the lifespan, we compared the expression profiles of each cell subset among different age groups. The differentially expressed genes (DEGs) among different age groups were identified in each subset. Then, the 31 PBMC subsets were ranked based on the number of DEGs (Figure 2A). The top ten cell subsets (DEG numbers > 300) belonged to lymphoid lineage cell subsets (i.e., T, NK and B cell types), indicating that these cell subsets were more susceptible to age. By contrast, not any myeloid lineages (i.e., monocytes, DCs and Mega) ranked the top ten cell subsets with DEG numbers > 300 (Figure 2A). Although eight of the top ten cell subsets belonged to T cells, significant difference in the number of DEGs was observed among different T cell subsets. Among T cell subsets, CD8_Trm_ZNF683 cells expressed the highest DEG numbers with 606 genes, however CD4_Treg_FOXP3 cells expressed the lowest DEG number with 98 genes (Figure 2A). Likewise, among NK cell subsets, NK_CD56^low^_FCGR3A cells expressed the highest DEG numbers with 460 genes, whereas NK_CD56^low^_TKTL1 cells expressed the lowest DEG number with 37 genes (Figure 2A). To gain an overview of functional dynamics of PBMCs across the life course, the DEGs in each subset were subjected to functional enrichment analysis. These DEGs were enriched in the biological processes related to antigen processing/presentation (in 29 cell subsets), interferon-gamma (IFN-γ) responses pathway (in 28 cell subsets), lymphocyte activation (in 27 cell subsets), ATP metabolic process (in 20 cell subsets), NF-κB signaling (in 18 cell subsets), and telomere maintenance (in 16 cell subsets) (Figure 2B).

**Figure 2.**
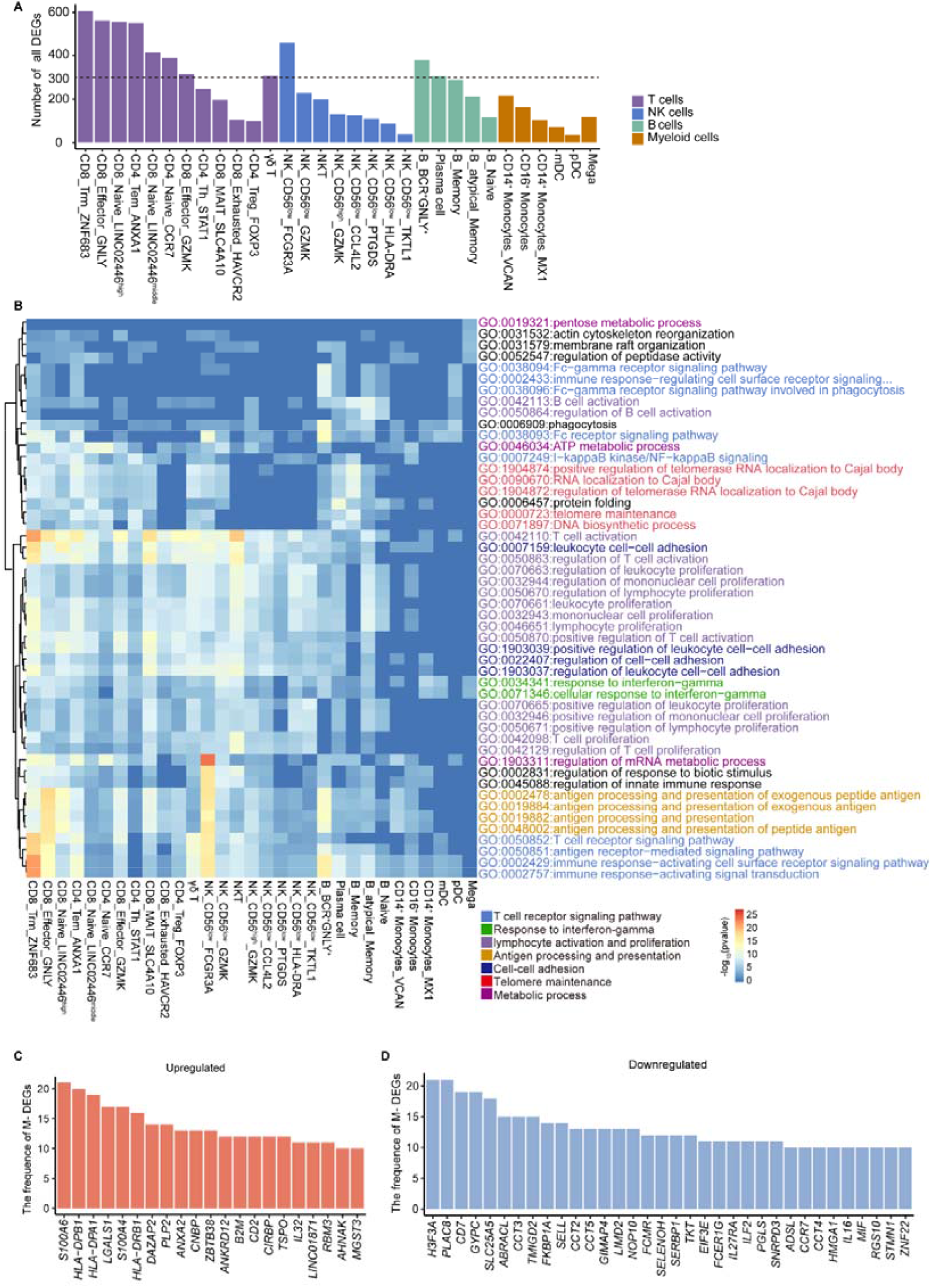
Global transcriptomic dynamics of peripheral immune cells throughout lifespan. (A) Bar plots showing the number of DEGs in all cell subsets. (B) Functional enrichment analysis of DEGs in all cell subsets. (C) Bar plots showing the frequencies of upregulated M-DEGs observed in all cell subsets and the frequency is greater than or equal to 10. (D) Bar plots showing the frequencies of downregulated M-DEGs observed in all cell subsets and the frequency is greater than or equal to 10.

Next, we focused on age-related genes that were monotonically correlated to age [named monotonical DEGs (M-DEGs); Upregulated or Downregulated] (details in Supplementary Methods). Detailed M-DEGs in each cell subset are listed in Table S3. Then, 31 PBMC subsets were ranked based on the number of M-DEGs (Figures S3A and S3B). All the top ten cell subsets (M-DEGs numbers > 100) belonged to lymphoid lineage cell subsets (i.e., T, NK and B cell types), implied that these lymphoid lineage cell subsets were susceptible to senescence. The top ten cell subsets included eight T cell subsets (CD8_Trm_ZNF683, CD4_Naive_CCR7 and CD8_Naive_LINC02446^high^, CD8_Naive_LINC02446^middle^, CD4_Tem_ANXA1, CD4_Th_STAT1, CD8_Effector_GZMK and CD8_Effector_GNLY cells), one NK cell subset (NK_CD56low_FCGR3A cells) and one B cell subset (B_Memory cells). The top three cell subsets with the largest number of M-DEGs were T cell subsets: CD8_Trm_ZNF683, CD4_Naive_CCR7 and CD8_Naive_LINC02446^high^ cells (Figures S3A and S3B). Functional enrichment analysis showed that the upregulated and downregulated M-DEGs were enriched in different biological processes. For instance, upregulated M-DEGs were enriched in NF-κB signaling and IFN-γ responses pathways (Figure S3C), while downregulated M-DEGs were enriched in telomere maintenance and proton transmembrane transport pathways (Figure S3E). To determine the common M-DEGs shared by multiple cell subsets, we ranked all M-DEGs based on the number of cell subsets in which they were involved. Results showed that 21 M-DEGs were upregulated in ≥ 10 cell subsets (Figure 2C), and 34 M-DEGs were downregulated in ≥10 cell subsets (Figure 2D). For example, the expression of genes involved in the IFN-γ signaling pathway (i.e., *HLA-DRB1, HLA-DPA1, HLA-DPB1* and *B2M*) was significantly upregulated in more than 10 cell subsets (Figures 2C and S3D). By contrast, the expression of genes involved in maintaining telomere length (i.e., *CCT3, CCT2* and *CCT5*) and nicotinamide adenine dinucleotide phosphate (NADP) metabolic process (i.e., *TKT* and *PGLS*) was significantly downregulated with age in more than 10 cell subsets (Figures 2D and S3F). These common M-DEGs across different cell subsets suggested that these genes may represent important immune biomarkers across the lifespan. Together, these results suggest that the functions of T cell subsets were more susceptible to the senescence among all PBMCs.

### Different aging patterns in naïve CD4^+^ T and naïve CD8^+^ T cells

A key feature of age-related immune erosion is the loss of naïve T cells (Goronzy and Weyand, 2019; Nikolich-Žugich, 2018). Proportion analysis of T cell subsets in total T cells showed that the proportions of all the three naïve T cell subsets [including two types of naïve CD8^+^ T cells (CD8_Naïve_LINC02446^high^ and CD8_Naïve_LINC02446^middle^) and one type of naïve CD4^+^ T cells (i.e.,CD4_Naïve_CCR7)] exhibited a decreased trend with age (Figure 1D and S2C), which was consistent with earlier flow cytometry results (van den Broek et al., 2018). We further compared the dynamic expression patterns of M-DEGs in naïve CD4^+^ T and naïve CD8^+^ T cells. The most of upregulated M-DEGs expression in naïve CD8^+^ T cells was sharply increased with age after 50 years old (i.e., 101 of 137 upregulated M-DEGs in CD8_Naïve_LINC02446^high^ cells, 69 of 115 upregulated M-DEGs in CD8_Naïve_LINC02446^middle^ cells) (Figures 3A, S4A and S4C), while the expression of all 99 upregulated M-DEGs in naïve CD4^+^ T cells (i.e., CD4_Naïve_CCR7) exhibited a progressive increase with age (Figure 3B). By contrast, no difference was observed in dynamic expression patterns of downregulated M-DEGs in naïve CD4^+^ T and naïve CD8^+^ T cells (Figures S4B and S4C). Moreover, functional enrichment analysis showed that the upregulated M-DEGs in naïve CD4^+^ T and naïve CD8^+^ T cells were significantly enriched in the biological processes related to inflammatory pathways, including “positive regulation of I-kappaB kinase/NF-kappaB signaling” and “response to interferon gamma” pathways (Figure S3C). We then analyzed the expression of these two inflammatory pathways in naïve CD4^+^ T and naïve CD8^+^ T cells across the whole lifespan. The results showed that the expression of these two inflammatory pathways in naïve CD8^+^ T cells (CD8_Naïve_LINC02446^high^ and CD8_Naïve_LINC02446^middle^) were sharply upregulated after 50 years old (Figures 3C, S4D and S4E), while displayed a gradual increase in naïve CD4^+^ T cells (CD4_Naïve_CCR7) (Figures 3C and S4D). Consistently, the variation of TCR diversity and “TCR V(D)J recombination” pathway expression was corelated with the abovementioned functional signatures in naïve CD4^+^ T and CD8^+^ T cells. Both the TCR diversity (Figures 3D and S4F) and “TCR V(D)J recombination” pathway expression (Figures 3E and S4G) in naïve CD8^+^ T cells (CD8_Naïve_LINC02446^high^ and CD8_Naïve_LINC02446^middle^) displayed an abrupt decrease after 50 years old, while those in naïve CD4^+^ T cells (CD4_Naïve_CCR7) showed a gradual decrease with age (Figures 3D and 3E). These results revealed the distinct aging-associated transcriptional patterns and repertoire dynamics in naïve CD4^+^ T and naïve CD8^+^ T cells.

**Figure 3.**
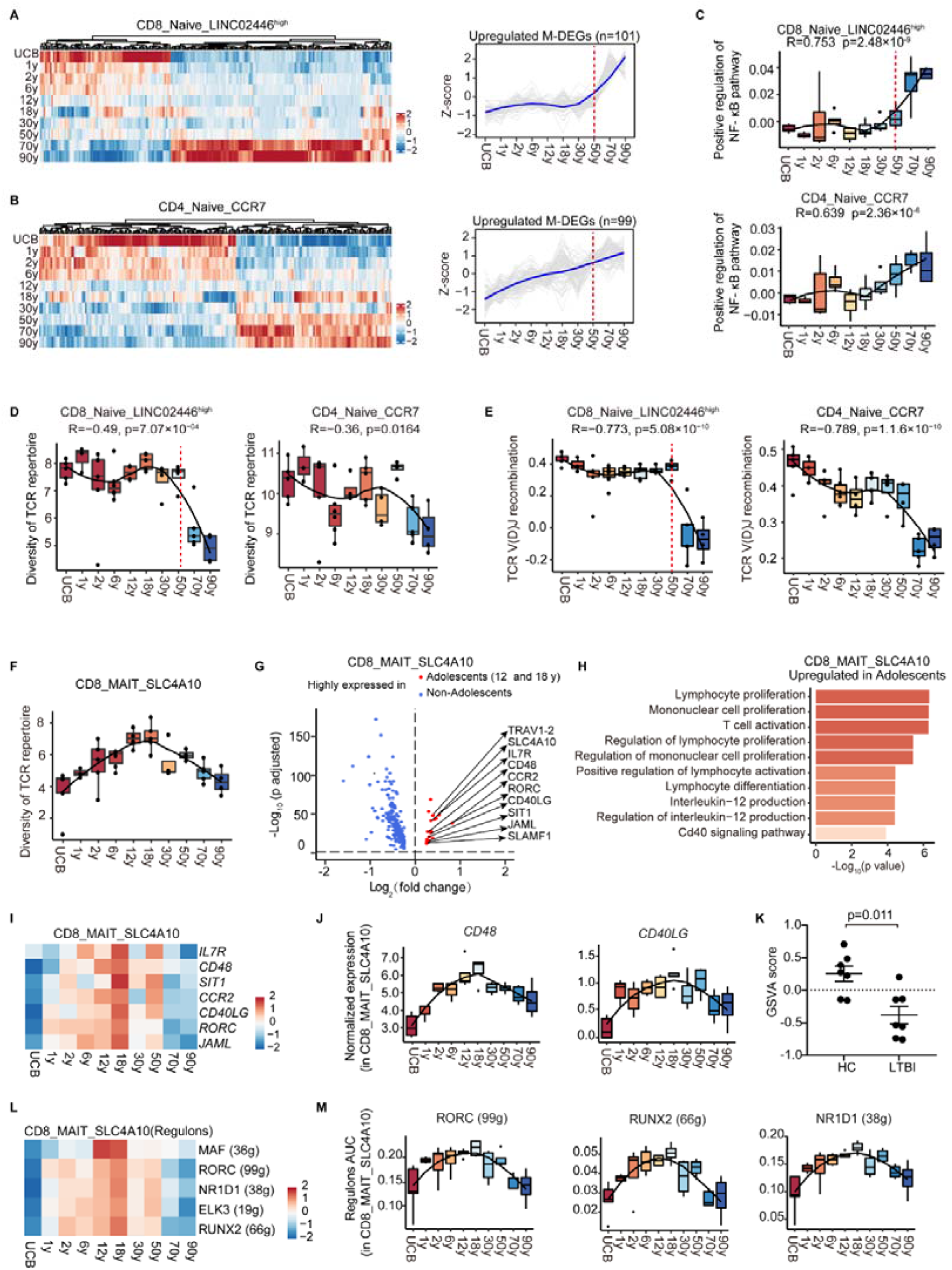
Aging features of naive T cells and CD8^+^ MAIT cells. (A) Heatmap showing the expression of M-DEGs in CD8_Naive_LINC02446^high^ cells across ten age groups (left panel). Expression trajectories of upregulated M-DEGs in CD8_Naive_LINC02446^high^ cells (right panel). Data are shown in terms of z-score of gene expression. Each grey line represents expression trajectory of each gene. Bule line represents the average expression trajectory. (B) Heatmap showing the expression of M-DEGs in CD4_Naive_CCR7 cells across ten age groups (left panel). Expression trajectories of upregulated M-DEGs in CD4_Naive_CCR7 cells (right panel). Data are shown in terms of z-score of gene expression. Each grey line represents expression trajectory of each gene. Bule line represents the average expression trajectory. (C) Box plots showing the expression of positive regulation of NF-κB signaling pathway in CD8_Naive_LINC02446^high^ and CD4_Naive_CCR7 cells across ten age groups. The black line indicates loess regression. Spearman correlation coefficient R and p value are indicated. A p-value□ <□ 0.05 was considered statistically significant. (D) Box plots showing the diversity of TCR repertoire in CD8_Naive_LINC02446^high^ (left) and CD4_Naive_CCR7 (right) cells across ten age groups. All box plots show median, 25th and 75th percentiles, and whiskers extending to maximum and minimum data points, with outliers beyond. The black line indicates loess regression. Spearman correlation coefficient R and p values are indicated. A p-value□ <□ 0.05 was considered statistically significant. (E) Box plots showing the expression of TCR V(D)J recombination in CD8_Naive_LINC02446^high^ (left) and CD4_Naive_CCR7 (right) cells across ten age groups. All box plots show median, 25th and 75th percentiles, and whiskers extending to maximum and minimum data points, with outliers beyond. The black line indicates loess regression. Spearman correlation coefficient R and p values are indicated. A p-value□ <□ 0.05 was considered statistically significant. (F) Diversity of TCR repertoire in CD8_MAIT_SLC4A10 cells across ten age groups. All box plots show median, 25th and 75th percentiles, and whiskers extending to maximum and minimum data points, with outliers beyond. The black lines indicate loess regression. Spearman correlation coefficient R and p values are indicated. (G) Volcano plot showing the DEGs in CD8_MAIT_SLC4A10 cells in adolescents compared with those in non-adolescents. Example genes are labeled with gene names. Red, upregulated in adolescents [Log_2_ (fold change) ≥ 0.25, adjusted p value < 0.05]; blue, upregulated in in non-adolescents [Log_2_ (fold change) ≤-0.25, adjusted p value < 0.05]. (H) Functional enrichment analysis of upregulated DEGs in CD8_MAIT_SLC4A10 cells in adolescents compared with those in non-adolescents. (I) Heatmap showing the expression of T cell proliferation and activation related genes in CD8_MAIT_SLC4A10 cells across ten age groups. (J) Box plots showing the expression of *CD48* and *CD40LG* in CD8_MAIT_SLC4A10 cells across ten age groups. All box plots show median, 25th and 75th percentiles, and whiskers extending to maximum and minimum data points, with outliers beyond. The black line indicates loess regression. (K) Scatter plots of GSVA score for T cell proliferation and activation related gene set in MAIT cells derived from healthy individuals (HC) and latent tuberculosis infection (LTBI) patients. Scatter dot plots showing mean□ ±□ SEM and p value determined by two-sided unpaired Wilcoxon test. A p-value□ <□ 0.05 was considered statistically significant. (L) Heatmap showing the regulon area under the curve of inferred differentially expressed TF regulons in CD8_MAIT_SLC4A10 cells across ten age groups. Numbers between brackets represent the regulon counts for respective transcription factors. Red, high expression; blue, low expression. (M) Box plots showing the area under curve of RORC (99g), RUNX2(66g) and NR1D1(38g) in CD8_MAIT_SLC4A10 cells across ten age groups. All box plots show median, 25th and 75th percentiles, and whiskers extending to maximum and minimum data points, with outliers beyond. The black line indicates loess regression. Numbers between brackets represent the regulon counts for respective transcription factors.

### Activation of CD8^+^ MAIT cells in adolescents

MAIT cells are unique innate-like T cells that bridge innate and adaptive immunity (Toubal et al., 2019). Accumulating evidence indicated that MAIT cells could rapidly produce cytokines and cytotoxic effectors via both TCR-dependent and -independent pathways (Legoux et al., 2020; Provine and Klenerman, 2020) in response to a broad range of bacteria and viruses, such as Mycobacterium tuberculosis (TB) (Ruibal et al., 2021) and Hepatitis B virus (Dias et al., 2019). Proportion analysis of T cell subsets showed that the proportion of CD8_MAIT_SLC4A10 cells in total T cells peaked at the adolescent stage (12 y and 18y groups) during the whole lifespan (Figure S2C), which was consistent with recent flow cytometry studies (Chen et al., 2019; Nicholas A et al., 2018). In addition, clonal analysis showed that the TCR diversity of CD8_MAIT_SLC4A10 cells peaked at the adolescent stage (12 y and 18y; Figure 3F), implying that CD8_MAIT_SLC4A10 cells from adolescents might more efficiently fend off pathogen invasion compared with other age groups. To further characterize the function of CD8_MAIT_SLC4A10 cells in adolescents, we compared the gene expression profiles of CD8_MAIT_SLC4A10 cells between adolescent (including 12 y and 18y) and other age groups. The upregulated DEGs in adolescent group were enriched in the biological processes related to T cell activation, lymphocyte proliferation and cytokine production (Figures 3G and 3H), while those downregulated DEGs in adolescents were enriched in the biological processes related to neutrophil activation and cell adhesion (Figure S4H). We further depicted the functional properties of CD8_MAIT_SLC4A10 cells in adolescents by GSEA. The GSEA results showed that the TGF-beta signaling, an immunosuppressive signaling, displayed a significant down-regulation in CD8_MAIT_SLC4A10 cells in adolescents (Figure S4I). To further determine the function of upregulated DEGs in adolescents, we downloaded the CD8^+^ MAIT cell transcriptome data of healthy individuals and latent tuberculosis infection patients (Pomaznoy et al., 2020). T cell proliferation and activation related genes including *IL7R, CD48, SIT1, CCR2, CD40LG, RORC* and *JAML*, highly expressed in adolescents (Figures 3I and 3J). Then, by Gene Set Variation Analysis (GSVA), we calculated the enrichment score of these T cell proliferation and activation related genes in MAIT cells derived from healthy individuals and latent tuberculosis infection patients. The GSVA enrichment score of healthy individuals was significantly higher than that of patients with latent tuberculosis infection (Figure 3K), possibly because of hypo-immune status in latent tuberculosis infection patients (Gong and Wu, 2021). According to the WHO 2021 Global Tuberculosis Report (https://www.who.int/), the TB incidence and prevalence in adolescents were lower than in adults (> 25 y), echoing our findings of enhanced MAIT cell functions in adolescents. Altogether, these results indicated that CD8_MAIT_SLC4A10 cells in adolescents were more resistant to pathogen invasion than those in other age groups.

To further investigate the transcriptional regulatory mechanism of CD8_MAIT_SLC4A10 cells in adolescents, we predicted the core transcription factors (TFs) using SCENIC (Aibar et al., 2017). We identified a set of TF regulons including RORC (99g), RUNX2 (66g), NR1D1 (38g), MAF (36g) and ELK3 (19g) by hierarchical clustering heatmap, whose regulon area under the curve values reached a peak in adolescents (Figures 3L, 3M and S4J). Moreover, four of five predicted TFs reached their expression peak in adolescents, including *RORC, RUNX2, NR1D1* and *ELK3* (Figure S4K). These TFs could directly regulate T-cell proliferation and activation-related genes highly expressed in adolescents, including *CD48, CD40LG, CCR2* and *IL7R* (Figure S6L). The transcription factor RORγt (encoded by *RORC*) has been reported to play crucial roles in MAIT cell maturation and effector functions by regulating IL-17 production (Guggino et al., 2017; Ivanov et al., 2006; Toubal *et al*., 2019). Together, these results implied that these TFs (including *RORC, RUNX2, NR1D1* and *ELK3*) might exert important roles in regulating the activation of CD8_MAIT_SLC4A10 cells during adolescence.

## Discussion

The human immune system dramatically changes across the lifespan (He and Sharpless, 2017; López-Otín and Kroemer, 2021). However, little is known about when and how the immune system changes across the entire human lifespan. Current study provided novel insights into the age-specific alteration in cell-type composition, transcription profiles, and immune repertoires of PBMCs across the entire human lifespan at single-cell resolution. Several novel contributions were made. First, we revealed that the functions of T cell subsets were the most susceptible to senescence among all PBMCs and were characterized by increased activation of NF-κB signaling and IFN-γ responses pathways, reduced telomere maintenance and energy metabolism. Second, scRNA-seq and immune repertoire sequencing disclosed unique transcriptional and clonal expansion features of T cell subsets [naïve CD4^+^ T and naïve CD8^+^ T cells exhibiting distinct aging-associated transcriptional patterns and repertoire dynamics; CD8_MAIT_SLC4A10 cells exhibiting peak abundance and clonal diversity in adolescents (12 y and 18 y groups)]. Overall, our dataset provides temporal resolution of immune cell transcriptomes across the entire human lifespan, and facilitates in-depth exploration of the immune system at specific ages.

The immune system undergoes dramatic changes beyond adolescence and sexual maturity (Vallejo, 2011). We found that the immune maturation during adolescence was characterized by the expansion of MAIT cells with higher diversity of TCR compared with other age groups. A previous study demonstrated that commensal bacteria mediated the MAIT cell maturation and its recirculation into thymus (Legoux et al., 2019), hinting a critical role of MAIT cells in TCR and immune system maturation. During adolescence, the dramatic changes of hormones lead to the microenvironment alteration of mucosal membrane and skin, and the variation of commensal bacterial species and numbers (Kundu et al., 2017), which regulate the proliferation of MAIT cells. MAIT cells contribute to host defense against infection (Toubal *et al*., 2019) including tuberculosis (Malka-Ruimy et al., 2019) and influenza (van Wilgenburg et al., 2018). According to the WHO 2021 Global Tuberculosis Report, the TB incidence and prevalence in adolescents were lower than in adults (> 25 y). However, until now, understanding of peripheral circulating MAIT cells in healthy individuals across the life course is still lacking. To our knowledge, the present findings for the first time delineated functional significance of MAIT cells in adolescence through the transcriptional profiling of this cell type across the lifespan.

The immune function declines with aging, termed “immunosenescence” (Nikolich-Zugich, 2018). However, different types of immune cells display differential susceptibility to aging. Our results revealed that the functions of T cell subsets were the most susceptible to senescence among all PBMCs, which were in a good agreement with the mice aging study by Matthew J et al. (Yousefzadeh *et al*., 2021). Moreover, functional enrichment analysis of cell type-specific age-related genes suggested that the elevation of pro-inflammatory signaling and the attenuation of signaling related to telomere maintenance and energy metabolism cooperatively contributed to immunosenescence and inflammaging. Furthermore, we discovered that naïve CD4^+^ and naïve CD8^+^ T cells exhibited distinct aging patterns. Both naïve CD4^+^ T and naïve CD8^+^ T cells arise from the thymus, but the behavior of naive T cells is different in CD4^+^ and CD8^+^ compartments. The expression of age-related genes and TCR diversity in naive CD8^+^ T cells exhibited sharp variations after 50 years old. By contrast, naïve CD4^+^ T cells showed progressive changes in expression of age-related genes and TCR diversity across the lifespan. Although the collapse of TCR diversity in naïve T cells has been considered a strong predictor of poor health in old people (Qi et al., 2014), the variation of TCR diversity with age in naïve CD4^+^ T and naïve CD8^+^ T cells is not very clear. Even though previous studies have reported that TCR diversity of naïve T cells displayed age-associated decline by bulk RNA sequencing (Britanova et al., 2014; Britanova et al., 2016), we depicted the different dynamics of TCR diversity across the lifespan in naïve CD4^+^ T and naïve CD8^+^ T cells for the first time. These results implied that the underlying mechanisms of aging in naïve CD4^+^ T and naïve CD8^+^ T cells are not identical, and provided valuable insights into the dysfunction of naïve T cells with age.

In conclusion, through our study, significant progresses have been made in understanding the age-related cellular and molecular variations in PBMCs during the development, maturation and aging of the immune system with greater details that were not appreciated previously. This dataset provides a valuable resource on circulating immune cells of different ages (available at https://pu-lab.sjtu.edu.cn/shiny for quick browsing), allowing for systematic elucidation of age-specific hallmarks of circulating immune cells across the lifespan, and serving as a reference for PBMCs dynamics during development, maturation and aging.

## ACKNOWLEDGMENTS

We thank the volunteers from the Pudong Natural Population-based Cohort for their contribution to the study. This work was supported by grants from the National Science Fund for Distinguished Young Scholars (81625002), the National Natural Science Foundation of China (81930007, 81470389, 81500221, 82071852, 82003013 and 81800307), the National High Level Talents Special Support Plan, the Shanghai Outstanding Academic Leaders Program (18XD1402400), Innovative research team of high-level local universities in Shanghai (SSMU-ZDCX20180200, SSMU-ZDCX20212100, SSMU-ZDCX20210601), Shanghai Municipal Education Commission Gao Feng Clinical Medicine Grant Support (20152209), Sponsored by Shanghai Sailing Program (21YF1445000), Shanghai Municipal Health Commission (20184Y0371) and the National Key R&D program of China (2021YFC2502300).

## AUTHOR CONTRIBUTIONS

J.P. conceived the study. R.Y.T., Z.L., J.F.S., A.C.Y., Y.P.G. and L.Y.S. recruited volunteers. R.Y.T., J.F.S., Y.G and Y.F.C. collected samples from donors. R.Y.T., J.F.S. and Y.Y. L. completed the experiments. R.H.L., R.Y.T., Y.C.Z, L.J.L., J.M.Z. and L.W. finished the data analysis. R.Y.T. and R.H.L drafted the initial version. Y.C.Z., Y.F.C. and L.J.L. proofed the manuscript. J.P., L.W., F.W., W.L, B.L, L.C, H.Y.C, and H.L.W provided guidance and constructive comments for the final version. All the authors read and approved the manuscript.

## STAR METHODS

### Participants and ethics

All human samples were obtained in accordance with protocols approved by the Institutional Review Board at Shanghai Jiao Tong University. All donors were enrolled with approval by the Ethics Committee of Ren Ji Hospital (KY2019-136), School of Medicine, Shanghai Jiao Tong University. After meeting inclusion criteria in accordance with the WHO classification and providing written informed consent in accordance with the criteria set by the Declaration of Helsinki (2013) (World Medical, 2013), 45 healthy volunteers from neonates to the elderly were recruited in the Pudong Natural Population-based Cohort, Shanghai, China. To avoid possible interference by other confounding factors, all participants were free of clinical symptoms including fever, cough, headache, sore throat, malaise, loss of smell, runny nose, abdominal pain, diarrhea, joint pain, wheezing or dyspnea and free of vaccination within three months before sampling. Meanwhile, the participants with hypertension, diabetes mellitus, autoimmune disease, cancer, renal or hepatic dysfunction, hematological disorders, pregnancy, lactation, steroid usage and smoking were excluded. They were assigned to 10 groups including group UCB standing for the neonates (umbilical cord blood, UCB); group 1 y, 2 y, 6 y standing for children aged 1, 2 and 6 years old respectively; group 12 y, 18 y standing for adolescents aged 12 and 18 years old; group 30 y, 50 y standing for adults aged 30 and 50 years old; group 70 y standing for septuagenarian aged 70 years old and group 90 y standing for nonagenarian aged ≥ 90 years old. Each group consisted of three to five individuals that included both males and females. Sample details are shown Table S1.

### Sample collection, preparation, and storage

Blood samples were collected from the peripheral vein of each donor and processed within 2 hours of collection except umbilical cord blood (UCB) obtained from placental cords. PBMCs were separated using Vacutainer CPT tubes (BD) according to the manufacture’s protocols (Corkum et al., 2015). The viability of PBMCs in each sample was confirmed to be >90% by Trypan Blue staining. Then, PBMCs were immediately used for scRNA library construction and sequencing or were cryopreserved at □ 80°C in 10% dimethylsulfoxide in fetal bovine serum.

### ScRNA-seq library construction and sequencing

To produce single-cell gel beads in emulsion (GEMs), cell suspensions were quantified with 400-600 living cells per microliter determined by a Count Star. Then, a Chromium single cell controller (10× Genomics) was used to prepare cell suspensions as GEMs with a Single Cell 5□ Library and Gel Bead kit (10× Genomics, 1000006) and Chromium Single Cell A Chip kit (10× Genomics, 120236) following the manufacturer’s protocols. About 15,000 cells were added to each channel and the recovered target cells was estimated to be about 8,000 cells. Single-cell RNA libraries were constructed the Chromium Single Cell 5□ v1.1 Reagent (10× Genomics, Human T Cell:1000005; Human B Cell: 1000016) and a unique sample index was included in each sequencing library. Finally, sequencing was performed with a sequencing depth of at least 100,000 reads per cell with a pair-end 150 bp (PE150) reading strategy using the Illumina platform (CapitalBio Technology, Beijing, China).

### Single-cell RNA sequencing data processing

All single-cell sequencing data were mapped to the human reference genome GRCh38 (10× Genomics, Version 3.0.0.) using STAR (version 2.5.1b). Unique molecular identifier (UMI) matrices were established for each sample using the CellRanger (10× Genomics, Version 3.0.2) pipeline and subjected to the standard processing pipeline that included quality control, normalization, dimension reduction and clustering by Seurat (Version 3.2.0). Specifically, cells with less than 400 expressed genes, or >10% mitochondrial content were excluded. Genes in the following analysis were confirmed with at least one feature count over three cells. To remove potential doublets, cells detected genes above 2500 are filtered out. Additionally, we applied Scrublet package (Wolock et al., 2019) in python to identify potential doublets. The doublet score for each single cell and the threshold based on the bimodal distribution was calculated using default parameters. The expected doublet rate was set to be 0.075, and cells predicted to be doublets or with doubletScore larger than 0.25 were filtered. Based on these measures, gene expression profiles were normalized for each cell under SCT-transform using the Seurat package. Subsequently, the ‘FindVariableFeatures’ function facilitated the selection of variable genes following the default operation. The gene expression matrixes were integrated together using “FindIntegrationAnchors” and the “IntegrateData” functions was adopted to correct the batch effects among samples by projecting all genes onto a low dimensional subspace based on canonical correlation analysis. The integrated data were saved for following analysis. Furthermore, a shared nearest neighbor graph was constructed based on the Euclidean distance in the low dimensional subspace spanned by the selected significant principal components. The “FindClusters” function was used to cluster these cells at an appropriate resolution. Finally, the “RunUMAP” function facilitated visualization of all these cells on the plane based on the two-dimensional uniform manifold approximation and projection (UMAP). Following the above pipeline, high-quality scRNA-seq data of ∼380k cells derived from 45 samples were remained.

### Cell types annotation and cluster markers identification

The “FindAllMarkers” function was used to find markers for each cluster. Clusters were then classified and annotated based on the signature gene expression of each cell subset.

All assigned cell subsets were based on the entire cluster instead of individual cells. Moreover, the variety within the same cluster was rectified by dominant prediction for the whole cluster. A full list of canonical and signature genes for each subset is deposited in Table S2.

### Identification of DEGs in peripheral immune cells

To investigate the effect of age on peripheral immune cells, we identified DEGs among ten age groups for each PBMC subset via following steps: 1) For each cell subset in one sample, the synthetic bulk expression value for each gene was generated by dividing the gene-specific UMI count with the total number of UMIs in the cell and multiplying by 10000 (“CP10K”). 2) Ribosome and mitochondrial genes were removed in all the samples. 3) The genes with expression more than 1 (CP10K > 1) in at least three samples were kept and analyzed by one-way analysis of variance (one-way ANOVA). 4) The genes with p value < 0.001 were considered to be DEGs. Age-related genes (monotonically correlated with age; M-DEGs) were further determined using Spearman Rho analysis (Rho ≥ 0.5, p < 0.01) based on all DEGs. CP10K were defined as:

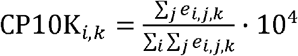

*e* is the number of UMIs of *i* gene defined by the unique molecular barcode of *j* cell in group *k*.

Next, to further investigate the expression profile of CD8_MAIT_SLC4A10 cells of adolescents, Seurat’s FindMarkers function (parameters: only.pos = F, min.pct = 0.25 logfc.threshold = 0.25, test = ‘wilcox’) was run comparing adolescents against non-adolescents. The DEGs were filtered using a minimum log_2_ (fold change) of 0.25 and a adjust p value of 0.05. The volcano plots of DEGs were performed by ggplot2 in R (version 3.6.0).

### Functional enrichment analysis and GSVA scoring

Functional enrichment analysis of DEGs were conducted using R package clusterProfiler (Yu et al., 2012) following the default parameters. R package REVIGO was applied to summarize Gene Ontology terms based on semantic similarity measures (Li et al., 2017; Supek et al., 2011). Gene set enrichment analysis (GSEA) was performed using the Java software GSEA 4.1.0 (Subramanian et al., 2005) based on the dataset ‘h.all.v7.1.sumbols.gmt [Hallmarks]’, ‘c2.cp.kegg.v7.1.symbols.gmt [Curated]’ and ‘c5.go.bp.v7.4.symbols.gmt’, collected from the Molecular Signatures Database (MSigDB) (Liberzon et al., 2011). The expression of signature genes in MAIT cells derived from healthy control (HC) and latent tuberculosis infection (LTBI) patients were quantified using GSVA package (Hänzelmann et al., 2013). The dataset on healthy individuals and latent tuberculosis infection patients came from GEO database (GSE132932) (Pomaznoy et al., 2020). Individuals GSVA scores were plotted as a scatter dot plot with the ggplots2 package. Next, a comparison of the GSVA enrichment score of healthy individuals and latent tuberculosis infection patients was performed using a two-sided unpaired Wilcoxon test, and a p-value < 0.05 was considered statistically significant.

### TCR and BCR repertoire sequencing and analysis

In addition to the single-cell transcriptome data, single-cell TCR and BCR repertoire libraries of all samples (10× Genomics) were prepared and profiled with paired-end sequencing (2□× □150□bp) on the Illumina HiSeq2500 platform (Illumina, San Diego, CA). The CellRanger V(D)J pipeline (10× Genomics, Version 3.0.2) was adopted to mark the TCR and BCR expression and clonal types. The TCR and BCR repertoire sequencing data were filtered with the following criteria: productive is “True”; high_confidence is “True”; umis ≥1; raw_consensus_id is not “None”. To identify the TCR clonotype for each T cell, only cells with at least one TCR α chain and one TCR β chain were used. For a given T cell, if there were two or more TCR α or TCR β chains assembled, the highest expressed (UMI or reads) TCR α or TCR β chains were regarded as the dominant TCR α or TCR β chain in the cell. Each unique TCR α(s)-TCR β(s) pair (CDR3 nucleotide sequences and rearranged VJ genes included) was defined as a clonotype. BCR clonotypes were identified in a similar manner to TCR clonotypes. Only cells with at least one productive heavy chain (IGH) and one productive light chain (IGL or IGK) were kept for further analysis. Each unique IGH-IGL/IGK pair was defined as a clonotype. For a given B cell, if there were two or more IGH or IGL/IGK assembled, the highest expressed (UMI or reads) IGH or IGL/IGK chain was defined as the dominant IGH or IGL/IGK chain in the cell. For CDR3 amino acid length analysis, filtered TCR α and β chains were separately calculated with “nchar”, “as.dat.frame”, “unlist”, “table”, “sum” and “round” functions. For the TCR clonal status analysis, “as.dat.frame”, “unlist” and “table” functions were used to calculate clonotype counts of filtered cells. Cells expressing the same clonotype were counted. The counts were classified into eight classes to identify the cellular clonal expansion status, i.e., >100, 51 to 100, 31 to 50, 21 to 30, 11 to 20, 6 to 10, 2 to 5, and a unique clone without expansion.

Moreover, based on the barcode mapping, clonal type information from CellRanger V(D)J was integrated into the meta data of the Seurat object for further analysis. To estimate TCR diversity, Shannon’s entropy was measured using the function ‘H’ of the philentropy package as follows:

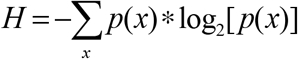

Here, the TCR among T with TCR identified is represented as p(x). Changes in clone diversity with age were fitted by loess regression. The Spearman correlation coefficient and significance were also calculated from the “cor.test” function in stats package. The p value<0.05 was considered statistically significant.

### Definition of cell state scores

Cellular modular scores were used to assess the cell state during the whole lifespan. The “AddModuleScore” function in Seurat was used to calculate the modular score of pathways with default settings. Response to interferon gamma (GO:0034341), Positive regulation of I-kappaB kinase/NF-kappaB (GO:0043123), V(D)J recombination (GO:0033151) were used to define the IFN-γ response, NK-κB signaling pathway, TCR V(D)J recombination. The cell scores for each pathway were calculated based on the average expression of genes from the abovementioned gene set. Expression signal changes with age were fitted by generalized loess regression. The Spearman correlation coefficient and significance were also calculated from the ‘cor.test’ function in stats package. A p-value < 0.05 was considered statistically significant.

### Transcription factor analysis

After arranging the input by the gene expression matrix, the SCENIC (Aibar *et al*., 2017) package in R facilitated evaluation of transcription factors among immune cell subsets following the default parameters. The pheatmap and ggplot2 packages in R were adopted to visualize the expression profile of transcription factors.

### Statistical analysis

Statistical analyses were performed using the R software package (version 3.6.0) and involved Shapiro-Wilk test, F-test, Levene’s test, one-way ANOVA, two-sided unpaired Wilcoxon test and Spearman’s correlation test. Data normality was determined by the Shapiro-Wilk test. The F-test was used to examine homogeneity of variance for two groups comparisons. The Levene’s test was used to examine homogeneity of variance for one-way ANOVA. We performed one-way ANOVA on cell subset proportions. When comparing GSVA enrichment score, two-sided unpaired Wilcoxon tests were used. Association between age and variables including cell subset proportions, gene expression and diversity of TCR was tested using a Spearman’s correlation test. The p value significance thresholds used for each result are described in the corresponding methods and figure legends.

### Data availability

The raw sequence data reported in this study will been deposited in the Genome Sequence Archive of the Beijing Institute of Genomics (BIG) Data Center, BIG, Chinese Academy of Sciences.

