## Supplemental figures for "Integrated single-cell sequencing analysis reveals peripheral immune landscape across the human lifespan": Supplemental_Biorxiv.docx

**Supplemental Figures and Figure legends**

**Supplemental Figure S1.**

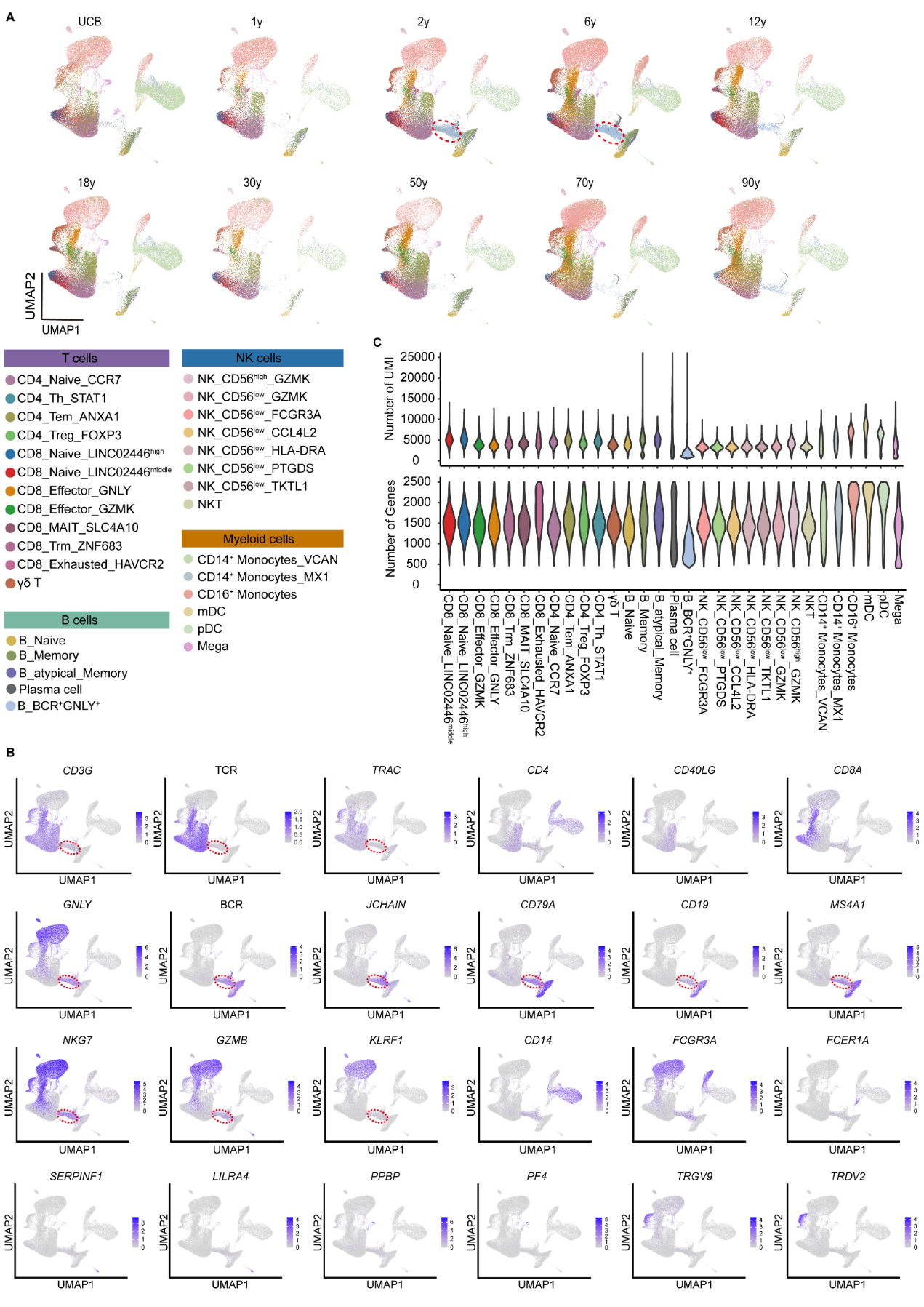

**Figure S1. Selected markers of cell subsets, UMAP projection of cell subsets and basic characteristics of the integrated dataset**
(A) UMAP charts of more than 380,000 single cells grouped by ten age groups. B_BCR^+^GNLY^+^ cells were circled with red dashed lines.

(B) Feature plots of selected signature genes for cell subsets. Expression levels were color-coded on the data, and the legend was labeled according to log scale. B_BCR^+^GNLY^+^ cells were circled with red dashed lines.

(C) Distribution of unique molecular identifier (UMI) counts per cell (top panel), gene counts per cell (bottom panel).

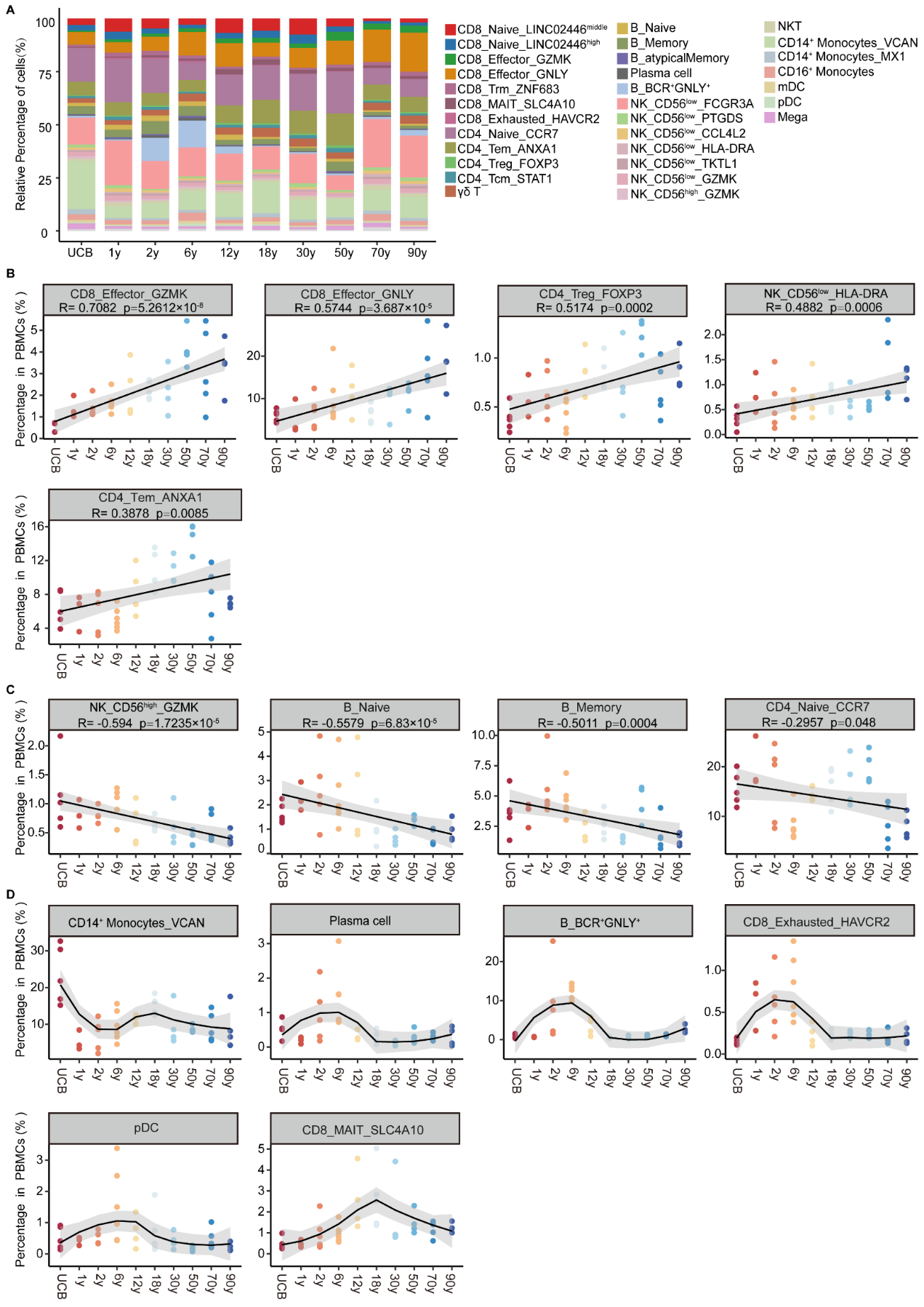

**Figure S2. Comparison of immune cell subsets based on unsort PBMC among ten age groups**

(A) Average proportion of 31 immune cell subsets in PBMC across ten age groups.

(B-D) Scatter plots showing the associations between age and cell subsets shown in Figure 1D. Spearman correlation coefficient R and p values are indicated. A p-value<0.05 was considered statistically significant.

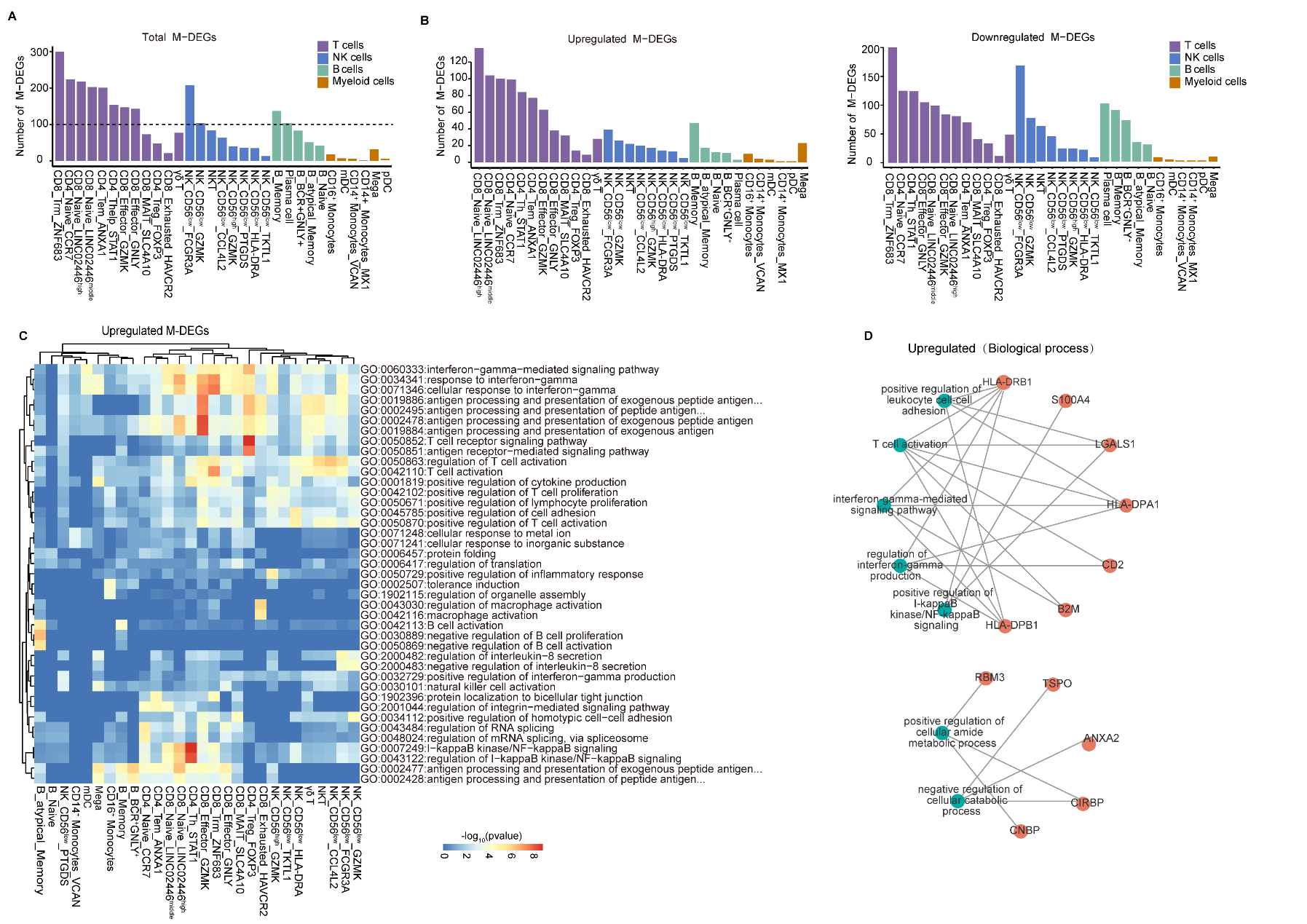

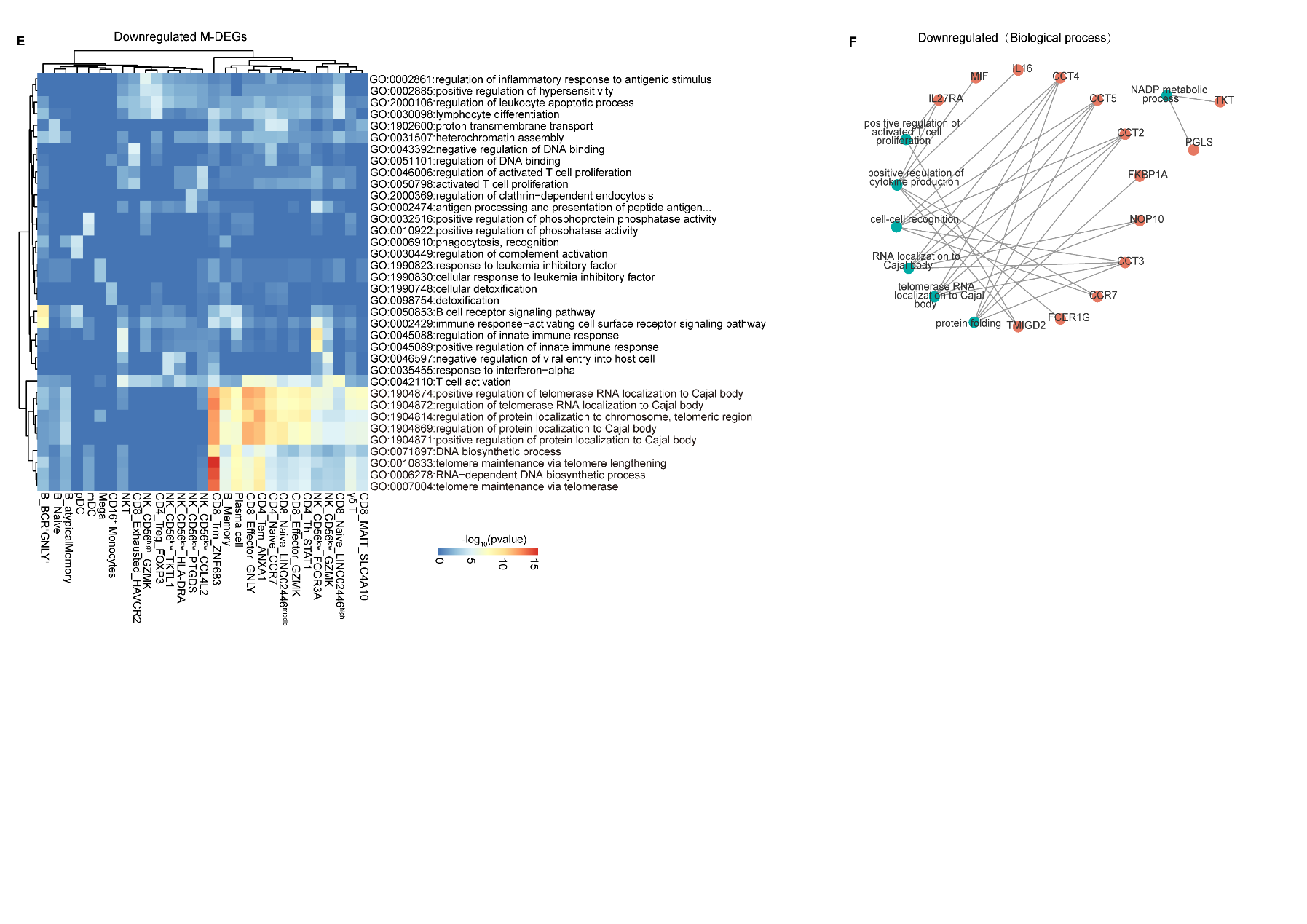

**Figure S3. Functional enrichment analysis of differentially expressed genes**

(A) Bar plots showing the numbers of total M-DEGs in all cell subsets.

(B) Bar plots showing the numbers of upregulated (left) and downregulated (right) M-DEGs in all cell subsets.

(C) Functional enrichment analysis of upregulated M-DEGs in all cell subsets.

(D) Functional enrichment network of the M-DEGs in (Figure 2C). The green nodes denote biological process terms; The orange red nodes denote the M-DEGs of these biological process terms. The size of green node represents the number of associated M-DEGs.

(E) Functional enrichment analysis of downregulated M-DEGs in all cell subsets.

(F) Functional enrichment network of the M-DEGs in (Figure 2D). The green nodes denote biological process terms; The orange red nodes denote the M-DEGs of these biological process terms. The size of green node represents the number of associated M-DEGs.

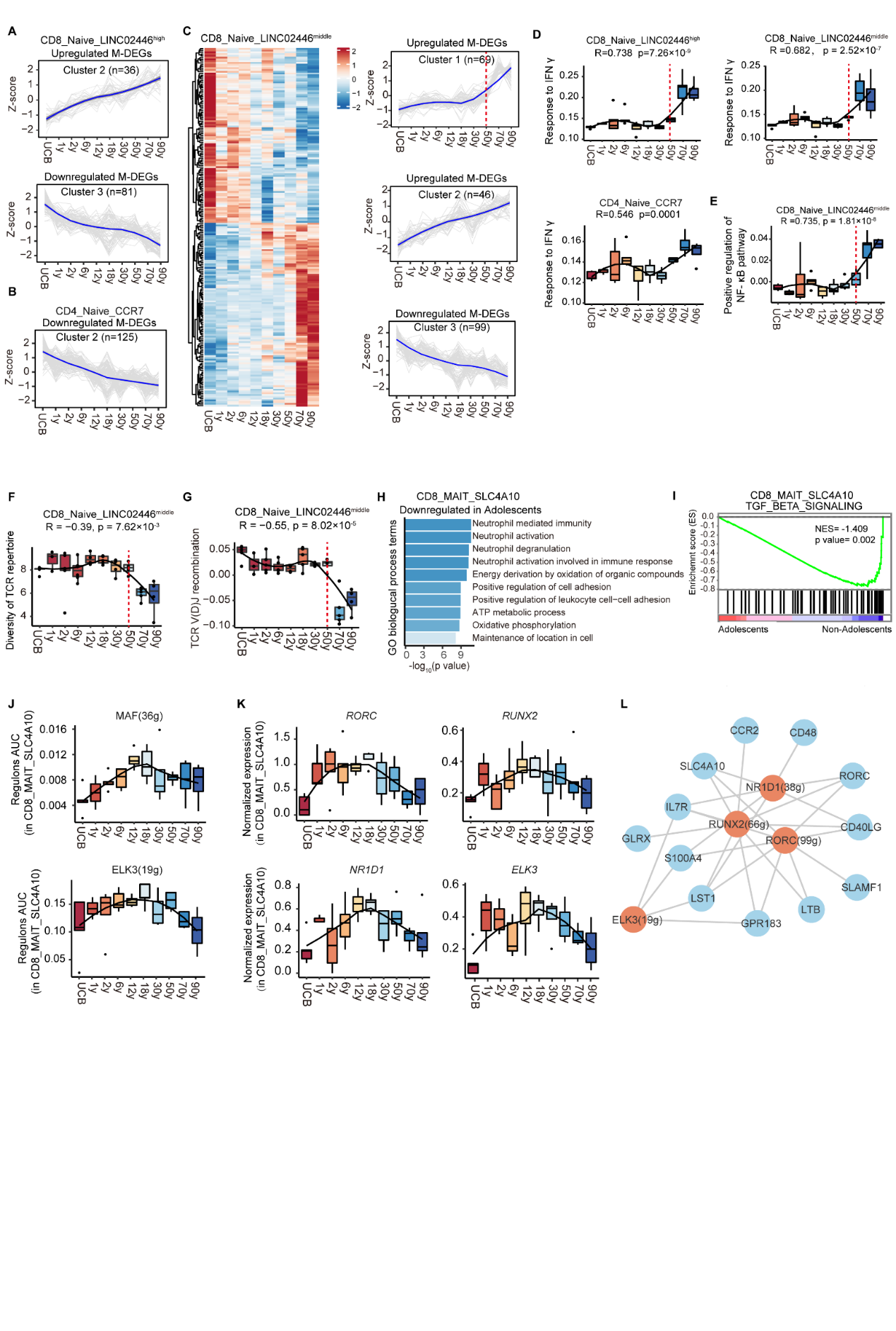

**Figure S4. Features of naïve T and CD8 MAIT cells**

(A) Expression trajectories of upregulated and downregulated M-DEGs in CD8_Naive_LINC02446^high^ cells. Data are shown in terms of z-score of gene expression. Each grey line denotes expression trajectory of each gene. Bule line denotes the average expression trajectory.

(B) Expression trajectories of upregulated M-DEGs in CD4_Naive_CCR7 cells. Data are shown in terms of z-score of gene expression. Each grey line denotes expression trajectory of each gene. Bule line denotes the average expression trajectory.

(C) Heatmap showing the expression of M-DEGs in CD8_Naive_LINC02446^middle^ cells across ten age groups (left panel). Expression trajectories of M-DEGs in CD8_Naive_LINC02446 ^middle^ cells (right panel). Data are shown in terms of z-score of gene expression. Each grey line represents expression trajectory of each gene. Bule line represents the average expression trajectory.

(D) Box plots showing the expression of response to interferon gamma pathway in CD8_Naive_LINC02446^high^, CD8_Naive_LINC02446^middle^ and CD4_Naive_CCR7 cells across ten age groups. The black line indicates loess regression. Spearman correlation coefficient R and p values are indicated. A p-value<0.05 was considered statistically significant.

(E) Box plots showing the expression of positive regulation of NF-κB signaling pathway in CD8_Naive_LINC02446^middle^ cells across ten age groups. The black line indicates loess regression. Spearman correlation coefficient R and p value are indicated. A p-value < 0.05 was considered statistically significant.

(F) Diversity of TCR repertoire in CD8_Naive_LINC02446^middle^ cells across ten age groups. All box plots show median, 25th and 75th percentiles, and whiskers extending to maximum and minimum data points, with outliers beyond. The black lines indicate loess regression. Spearman correlation coefficient R and p values are indicated. A p-value<0.05 was considered statistically significant.

(G) Expression of TCR V(D)J recombination pathway in CD8_Naive_LINC02446^middle^ cells across ten age groups. All box plots show median, 25th and 75th percentiles, and whiskers extending to maximum and minimum data points, with outliers beyond. The black lines indicate loess regression. Spearman correlation coefficient R and p values are indicated. A p-value<0.05 was considered statistically significant.

(H) Functional enrichment of downregulated DEGs in CD8_MAIT_SLC4A10 cells in adolescents compared with those in non-adolescents.

(I) GSEA enrichment plots showing one upregulated gene set downregulated gene set in CD8_MAIT_SLC4A10 cells in adolescents compared with those in non-adolescents. NES, normalized enrichment score. A p-value < 0.05 was statistically significant.

(J) Box plots showing the area under curve of MAF (36g) and ELK3(19g) in CD8_MAIT_SLC4A10 cells across ten age groups. Numbers between brackets represent the regulon counts for respective transcription factors. All box plots show median, 25th and 75th percentiles, and whiskers extending to maximum and minimum data points, with outliers beyond. The black line indicates loess regression.

(K) Box plots showing the expression of *RORC*, *RUNX2*, *NR1D1* and *ELK3* in CD8_MAIT_SLC4A10 cells across ten age groups. All box plots show median, 25th and 75th percentiles, and whiskers extending to maximum and minimum data points, with outliers beyond. The black line indicates loess regression.

(L) Network visualization of TF regulons and their target genes. Numbers between brackets represent the regulon counts for respective transcription factors. The orange red nodes annotate TF regulons; the light blue nodes denote the target genes of these TF regulons.

**Supplementary Tables**

**Table S1. Detailed characteristics of 45 healthy individuals.**

**Table S2. Marker genes of cell subsets.**

**Table S3. Monotonous genes associated with age.**
